# HNRNPM promotes protein synthesis via an interaction with stem-loop 5 in the SARS-CoV-2 5’-UTR

**DOI:** 10.64898/2026.10.05.756635

**Authors:** Jasmin Golagha, Maren Wieja, Nadine Biedenkopf, Julia E. Weigand

## Abstract

The 5’-untranslated region (5’-UTR) of severe acute respiratory syndrome coronavirus 2 (SARS-CoV-2) is highly structured and comprises five conserved stem-loops that serve as *cis*-acting regulatory elements. During infection, host proteins are recruited to these elements and regulate multiple steps during the viral life cycle. Stem-loop 5 (SL5) is highly conserved among different alpha- and betacoronaviruses and includes the start codon of non-structural protein 1 (nsp1). However, the function of SL5 in viral replication and its interacting proteins remain to be determined. Here, we identified heterogeneous nuclear ribonucleoprotein M (HNRNPM) as a host protein functionally interacting with SL5. HNRNPM translocates into the cytoplasm during SARS-CoV-2 infection and promotes protein synthesis from the viral 5’-UTR. Mutations in the lower stem of SL5 impair HNRNPM binding and reduce protein synthesis. Notably, the interaction between HNRNPM and SL5 is conserved among different alpha- and betacoronaviruses. Our findings indicate that HNRNPM is an important host factor that regulates viral translation to promote SARS-CoV-2 replication.

**IMPORTANCE:** SARS-CoV-2 causes the coronavirus disease 2019 (COVID-19), which has demonstrated the pandemic potential of coronavirus infection and the need for effective antiviral treatment. The 5’-UTR of SARS-CoV-2 contains essential structured RNA elements, yet the function of many of these elements, including SL5, remain poorly understood. Our findings provide the first indication that SL5 can serve as a platform for modulation of viral protein synthesis by host factors and offer the first mechanistic understanding into the function of the proviral factor HNRNPM in SARS-CoV-2 infection. This regulation provides new insight into how SARS-CoV-2 employs host factors and could provide a strategy to limit viral replication.

## INTRODUCTION

SARS-CoV-2 is a positive-sense, single-stranded RNA virus with an approximately 30 kb genome (1). It belongs to the family *Coronaviridae*, which is classified into four genera. Alphacoronaviruses (e.g., HCoV-229E and HCoV-NL63) and betacoronaviruses (e.g., SARS-CoV-2, MERS-CoV and HCoV-OC43) infect mammals, and include all known human coronaviruses, whereas gammacoronaviruses and deltacoronaviruses most commonly infect birds (2). During infection, SARS-CoV-2 releases its 5’-capped and 3’-polyadenylated genome into the host cell cytoplasm (3). Subsequently, the host ribosomes translate the open reading frames (ORFs) 1a and 1b to synthesize the polyprotein pp1a and the frameshift-dependent pp1ab. These polyproteins are cleaved into 16 nonstructural proteins (nsps) and assemble the replication-transcription complex (RTC). Synthesis of both polyproteins depend on translation initiation at the first genomic AUG, which is embedded in a highly structured region at the genomic 5’-end.

The 5’-UTR of SARS-CoV-2 encompasses nucleotides (nts) 1-265 and contains five stem-loops (SLs) that function as essential *cis*-regulatory elements controlling RNA translation, transcription and replication (4). Stem-loop 1 (SL1) enables viral transcripts to escape nsp1-mediated translational repression and RNA degradation (5–8). SL5 comprises nucleotides 150-294, including the start codon and 29 nt of the ORF of nsp1, the first encoded non-structural protein. SL5 folds into a four-way junction with a stem part and three sub-SLs (SL5a, SL5b, and SL5c) (4). Its three-dimensional structure of is broadly conserved across alpha- and betacoronaviruses and characterized by a T-shaped conformation (9, 10). SL5a and SL5b contain the highly conserved apical loop motif 5’-UUYCGU-3’ (11). In contrast, the apical loop of SL5c shows greater sequence variability. Alphacoronaviruses mostly exhibit the same apical loops in SL5a-c, while in SARS-CoV-2, SL5c harbors the tetra-nucleotide loop sequence 5’-GAAA-3’. The SL5 stem part is formed by base pairing between the nts 150-181 and nts 264-294 and is interrupted by several single-stranded regions. The stability and local dynamics of the SL5 stem are highly temperature dependent (12).

So far, the function of the SL5 structure is unclear. The conserved SL5a and SL5b loop motifs have been considered to act as packaging signal (4, 11). However, initial virus-like particle assays identified PS9, a sequence within the nsp15/16-coding region (13), while a later study pointed towards a pseudoknot structure in the nsp12-coding region (14). Just recently, the SARS-Cov-2 SL5 has been suggested to be part of an internal ribosome entry site (IRES), particularly aiding viral translation under hypoxic conditions (15). Here, the apical loops of SL5a and SL5b have been suggested to act a recognition site for unknown IRES *trans*-acting factors (ITAFs).

Generally, the function of *cis*-acting RNA elements is mediated through interactions with *trans*-acting factors, including viral and host RNA-binding proteins (RBPs). Several SARS-CoV-2 RNA interactome studies have identified numerous host proteins that associate with the SARS-CoV-2 genome and influence viral replication (16–20). For example, it has been shown that the RNA-binding protein ZAP inhibits the SARS-CoV-2 frameshifting by binding the pseudoknot structure in the nsp12-coding region (17, 21). An early interactome study performed knockdown experiments targeting host RBPs to assess their function on viral replication (19). Depletion of multiple factors either upregulated or downregulated viral RNA levels, suggesting that many host RBPs retain pro- or antiviral functions. One of the RBPs with the highest proviral effect observed was HNRNPM.

Thus, characterizing the host proteins that bind to SL5 may reveal new opportunities to reduce viral replication by targeting the RNA or by interfering with proviral host proteins it recruits. Here, we investigate SARS-CoV-2 SL5 as a *cis*-regulatory element and establish its interaction with host RBPs. We perform RNA pulldown assays and identify HNRNPM as functional SL5 interactor. Further, we show that this RNA-protein interaction affects protein synthesis from the viral 5’-UTR.

## MATERIALS AND METHODS

### Plasmid construction

#### pHDV plasmids

DNA templates comprising the T7 promoter upstream of the different 5’-SL5 variants (SL5wt, loop mut, shuffled, antisense, 4-way, stem, stem mut, SL5 MERS-CoV, SL5 HCoV-229E) were prepared by hybridization and phosphorylation of complementary oligonucleotides (Supplementary Table S1). Inserts were cloned into the pHDV vector (22) via *Eco*RI and *Nco*I restriction sites.

#### pCMV-HNRNPM plasmids

Total RNA was extracted from A549 cells (DSMZ, ACC107). A549 cells were treated with TRIzol Reagent (Invitrogen) followed by chloroform extraction and isopropanol precipitation. 1 µg total RNA was incubated with Oligo d(T)20 primer and dNTPs for 5 min at 65°C and reverse transcribed for 10 min at 55°C using SuperScript IV reverse transcriptase (Invitrogen) followed by heat inactivation for 10 min at 80°C. RNA was removed with RNase H (NEB) for 20 min at 37°C. cDNA was used as template for PCR to amplify HNRNPM ΔNLS (Supplementary Table S2), while introducing *Bsa*I and *Xho*I restriction sites. The PCR product was purified with DNA Clean & Concentrator-5 (Zymo Research) and digested with BsaI and XhoI. A FLAG-tag was obtained by hybridization and phosphorylation of complementary oligonucleotides and was digested with MluI and BsaI. The pCMV-MS vector (23) was digested with MluI and XhoI. Digested DNA was gel-purified using the Monarch® DNA Gel Extraction Kit (NEB). FLAG and HNRNPM inserts were ligated into the pCMV-MS vector. A clone encoding for the HNRNPM isoform b (NM_031203) was selected. Full-length and mutated HNRNPM plasmids were obtained by site-directed mutagenesis PCR (Supplementary Table S2).

#### Dual luciferase reporter plasmids

Dual luciferase reporter plasmids were constructed according to (8), with NanoLuc luciferase (nluc) as reporter gene and firefly luciferase (fluc) as internal control. To facilitate seamless integration of coronaviral 5’-UTRs, two *BsaI* restriction sites were cloned upstream of the nluc ORF by site-directed mutagenesis PCR, resulting in the vector pDLNluc-GG. The 5’-UTR wt sequence of SARS-CoV-2 was obtained from pSP64_SCoV2_5’-UTR (12) via PCR and inserted into pDLNluc-GG using the *Bsa*I restriction sites. Mutants were generated by site directed mutagenesis PCR or inverse PCR. 5’-UTR sequences of MERS-CoV, MHV-A59, HCoV-OC43, HCoV-229E, HCoV-NL63 were ordered from GeneArt ThermoFisher and inserted into pDLNluc-GG via *Bsa*I restriction sites (Supplementary Table S3).

### *In vitro* transcription

pHDV plasmids were linearized with HindIII and purified by phenol-chloroform extraction and isopropanol precipitation. 100 µg linearized plasmid was *in vitro* transcribed overnight at 37°C with 5-10% DMSO, 20-30 mM magnesium acetate, 200 mM Tris-HCl pH 8.0, 20 mM DTT, 2 mM spermidine, 4 mM NTP each and 3% (v/v) T7 RNA polymerase (in-house purified). RNA was transcribed as an HDV ribozyme fusion to achieve 3’-end homogeneity. Purification was performed using 6% denaturing polyacrylamide gels. RNA length and integrity was verified on a control gel.

### Pulldown

#### 5’-RNA biotinylation

RNA was dephosphorylated with Antarctic Phosphatase (NEB) for 30 min at 37°C and subsequently phosphorylated with T4 polynucleotide kinase (NEB) and ATPγS (Biomol) for 30 min at 37°C. Phosphorylated RNA was then incubated with biotin long arm maleimide (vector laboratories) for 30 min at 65°C. Biotinylated RNA was purified by phenol-chloroform extraction and ethanol precipitation.

#### Cell lysate

A549 cells were detached using Trypsin-EDTA two days after seeding. Collected A549 cells were centrifuged for 10 min at 500 x g and washed with PBS. Cell pellet was resuspended with lysis buffer (10 mM Tris-HCl pH 7.4, 1.5 mM MgCl_2_, 150 mM KCl, 0.5 mM DTT, 0.5% Igepal, 100 U/mL RNase inhibitor, 1% protease inhibitor cocktail, 1% phosphatase inhibitor) and snap-freezed in liquid nitrogen. Lysates were centrifuged for 10 min at 16,000 x g at 4°C. Supernatant was collected and protein concentration was determined via Bradford assay.

#### Pulldown

50 µl Dynabeads M-280 Streptavidin (Invitrogen) were washed with RNA incubation buffer (10 mM Tris-HCl pH 7.4, 1.5 mM MgCl_2_, 150 mM KCl, 0.5 mM DTT, 0.05% Igepal) according to the manufacturer’s protocol. 100 pmol biotinylated RNA was incubated with the beads for 1 h at 4°C with rotation. Supernatant was discarded and beads were washed with RNA incubation buffer. 500 µg protein lysate mix [10 mM Tris-HCl pH 7.4, 1.5 mM MgCl_2_, 150 mM KCl, 0.5 mM DTT, 0.5% Igepal, 180 U/ml RNase inhibitor (Molox), 200 µg/mL Heparin, 200 µg/mL tRNA] was added to the RNA-bound beads and incubated for 2 h at 4°C with rotation. Beads were washed with RNA incubation buffer. Bound proteins were eluted with SDS loading buffer (15% 2-mercaptoethanol, 4% SDS, 50% glycerin, bromophenol blue) at 95°C for 10 min. Supernatant was analyzed via SDS-PAGE and western blot.

### Mass spectrometry

Coomassie stained gel-slices were cut out of an SDS-PAGE and destained by the addition of 200 µL destaining solution (50 mM NH_4_HCO_3_ in 50% MeCN) and incubation for 30 min. Supernatant was then discarded and destaining was repeated till the gel slice was completely destained. Subsequently the gel-slices were dried in a vacuum concentrator for ~15 min at 45°C. Trypsin (0.1 µg in 30µL 50 mM NH_4_HCO_3_) was added to each sample and samples were incubated at 37°C overnight. On the next day, 200 µL MeCN were added, and samples were incubated in an ultrasonic-bath for 30 min. The supernatant was transferred to fresh tubes and evaporated to dryness using a vacuum concentrator. The pellet was dissolved in 25 µL 5% MeCN with 0.1% formic acid.

The mass spectrometric analysis of the samples was performed using a timsTOF Pro mass spectrometer (Bruker Daltonics). A nanoElute HPLC system (Bruker Daltonics), equipped with an Aurora Ultimate C18 RP column (25cm x 75µm ID) filled with 1.7 µm beads (IonOpticks, Australia) was connected online to the mass spectrometer. 2 µL of the sample was injected directly on the separation column. Sample loading was performed at a constant pressure of 800 bar. Separation of the tryptic peptides was achieved at 60°C column temperature with the following gradient of water/0.1% formic acid (solvent A) and acetonitrile/0.1% formic acid (solvent B) at a flow rate of 400 nL/min: Linear increase from 2% B to 17% B within 18 min, followed by a linear gradient to 25% B within 9 min and linear increase to 37% solvent B in additional 3 min. Finally, B was increased to 95% within 10 min and hold at 95% for additional 10 min. The built-in “DDA PASEF-standard_1.1sec_cycletime” method developed by Bruker Daltonics was used for mass spectrometric measurement.

Raw Data was automatically processed within “Data Analysis” (Bruker Daltonics, Bremen) with the method “DA_nanoElute wo cal wo xml - PASEF.m” provided by Bruker. This method writes an MGF-File (Mascot Generic Format), which was subsequently searched against the uniprot human database and common contaminants using Proteome Discoverer 2.4 software (Thermo Fisher Scientific, Bremen) (Supplementary Tables S4 and S5).

### Infection of A549-hACE2-TMPRSS2

A549-hACE2-TMPRSS2 cells (InvivoGen) were seeded 24 h prior to infection at a density of 200,000 cells per well in a 6-well plate in DMEM (Gibco) supplemented with 10% FCS (Thermo Fisher Scientific), 100 U/mL penicillin (Gibco), 100 µg/mL streptomycin (Gibco), 2 mM L-glutamine (Thermo Fisher Scientific), 0.5 μg/mL puromycin (InvivoGen) and 300 μg/mL hygromycin (InvivoGen). Cells were infected with BavPat1/2020 isolate (European Virus Archive Global #026V-03883) at a multiplicity of infection (MOI) of 0.1 for 1 h at 37°C in DMEM supplemented with 0% FCS, 100 U/mL penicillin, 100 µg/mL streptomycin, 2 mM L-glutamine. The inoculum was replaced by DMEM supplemented with 10% FCS, 100 U/mL penicillin, 100 µg/mL streptomycin, 2 mM L-glutamine. All work with live SARS-CoV-2 was performed in the BSL-4 facility (Institute of Virology, Marburg University).

### Indirect immunofluorescence analysis

Indirect immunofluorescence analysis was performed as described in (24). Cells were fixed before infection as well as 6 h and 24 h post infection (p.i.) in 4% paraformaldehyde. Thereafter cells were treated with 0.1 M glycine before being permeabilized using 1% TritonX100 followed by incubation in blocking buffer (PBS with 2% BSA, 5% glycerin, 0.02% Tween20, 0.05% NaN_3_). Staining was performed using protein-specific antibodies for HNRNPM (Santa Cruz, sc-20002) and SARS-CoV-2 N (Sino Biological, 40143-R001). Nuclei were stained using 0.1 µg/ml DAPI (Sigma-Aldrich). Cover slips were fixed with FluorSave Reagent (Millipore Sigma). Images were obtained using a Leica DMI6000B (63x magnification, oil immersion objective). Image processing and analysis was performed using ImageJ/Fiji v1.54f. For the quantification of the mean intensity of cytoplasmic and nuclear HNRNPM signal, nuclei were outlined using the Particle Analyze tool, cell borders were outlined manually. Background subtraction was performed using the Rolling Ball Background Subtraction plugin. For each biological replicate 50 cells per condition were quantified.

### Transfection

Human A549 (DSMZ, ACC107) cells were cultured in DMEM (Gibco) supplemented with 10% fetal bovine serum (Sigma-Aldrich), 1% penicillin-streptomycin (Gibco) and 1% sodium pyruvate (Gibco) at 37°C and 5% CO_2_. Transient transfections were performed using Lipofectamine 3000 (Thermo Fisher Scientific) according to the manufacturer’s protocol.

#### Transfection for dual luciferase assays

A549 cells were seeded in a 96-well plate at a density of 15,000 cells per well. 10 ng pDLNluc plasmid were co-transfected either with 300 ng of a pCMV-HNRNPM variant or 300 ng peGFP (23) control plasmid 24 h after seeding.

#### Transfection for western blots

A549 cells were seeded in a 12-well plate at a density of 140,000 cells per well. 3 µg of a pCMV-HNRNPM variant or 3 µg of peGFP control plasmid were transfected 24 h after seeding.

#### Transfection for total RNA isolation

A549 cells were seeded in a 12-well plate at a density of 140,000 cells per well. 100 ng pDLNluc-GG plasmid were co-transfected either with 3 µg pCMV-HNRNPM variant or 3 µg peGFP control plasmid 24 h after seeding.

### Protein lysates

#### Subcellular fractioning

Nuclear and cytoplasmic extracts were collected using NE-PER Nuclear and Cytoplasmic Extraction Kit (Thermo Scientific) according to the manufacturer’s protocol. Protein samples from SARS-CoV-2 infected A549-hACE2-TMPRSS2 cells were collected in sample buffer (167 mM Tris HCl pH 6.8, 20% 2-mercaptoethanol, 8% SDS, 33.3% glycerol, bromophenol blue) before infection and 24 h post infection. Subcellular protein fractions from transfected A549 cells were collected 24 h after transfection and quantified using Bradford assay.

#### Total Protein Lysate

A549 cells were lysed with lysis buffer (137.5 mM NaCl, 10% glycerin, 20 mM Tris-HCl pH 8.0, 2 mM EDTA, 1% Igepal, 1% protease inhibitor cocktail) on ice for 20 min 24 h after transfection. Lysate was centrifuged for 15 min at 16,000 x g at 4°C. Supernatant was collected and protein concentration was measured via Bradford assay.

### Western blot

5 µg of total protein or subcellular protein fraction were separated by SDS-PAGE. Total protein was imaged as loading control on the ChemiDoc Imaging System (Bio-Rad). Proteins were transferred to a PVDF membrane using the Trans-Blot turbo transfer system (Bio-Rad) with the “Mixed MW” program. The membrane was incubated with primary antibody [anti-HNRNPM antibody (Santa Cruz, sc-20002), anti-Vinculin antibody (Sigma-Aldrich, V9131), anti-SARS-CoV-2 Nucleocapsid antibody (Sino Biological, 40143-R001)] in 2% milk overnight at 4°C or with HRP conjugated anti-FLAG antibody (proteintech, HRP-66008) in 5% milk for 2 h at room temperature. Secondary antibodies [HRP-conjugated anti-mouse IgG (Jackson Immunoresearch), HRP-conjugated anti-rabbit IgG (Jackson Immunoresearch)] were used for detection of primary antibodies without HRP conjugate. The protein-antibody-complex was detected with WesternBright Chemilumineszenz Substrat Sirius (Advansta) according to the manufacturer’s protocol and quantified with the Image Lab software (Bio-Rad).

### Dual luciferase assay

Readout was performed 24 h after transfection with the Nano-Glo Dual-Luciferase Reporter Assay System (Promega) according to the manufacturer’s protocol. Luminescence was measured using a TECAN plate reader. NanoLuc luciferase (nluc) activities were normalized to firefly luciferase (fluc) activities, which served as an internal control. Data are presented as RLU_nluc_/RLU_fluc_ ratios.

### RNA isolation and quantitative RT-PCR

Total RNA from A549 cells was extracted with TRIzol reagent (Invitrogen), purified with the RNeasy Mini Kit (Qiagen) and treated with DNase for 1 h at 37°C. RNA was purified again using the RNeasy Mini Kit. RNA integrity was checked on a 1% agarose gel. 300 ng of total RNA were annealed with random hexamers (Jena Bioscience) at 70°C for 5 min. RNA was reverse transcribed in a 20 µl reaction volume using GoScript reverse transcriptase (Promega). The reaction mix was incubated at 25°C for 5 min, followed by 42°C for 45 min and then heat inactivated at 70°C for 15 min. The resulting cDNA was diluted to a final volume of 60 µl. The qPCR reaction was prepared with 5 µl cDNA, Fast SYBR Green Master Mix (Thermo Fisher Scientific) and target-specific oligonucleotides (nluc: forward: 5’-CAGCCGGCTACAACCTGGAC-3’, reverse: 5’-AGCCCATTTTCACCGCTCAG-3’ (25); fluc: forward: 5’-GCTCAGCAAGGAGGTAGGTG-3’, reverse: 5’-TCTTACCGGTGTCCAAGTCC-3’ (23)) in a total reaction volume of 20 µl. nluc and fluc levels were quantified with the QuantStudio 3 Real-Time-PCR system (Thermo Fisher Scientific). Melting curve analysis confirmed product specificity. mRNA levels were calculated with 2^−ΔCt^ and normalized to the expression level of the control construct with eGFP co-transfection.

### Software

RNA drawings were created with RNA canvas (26). All statistical analyses were performed using GraphPad Prism v10.

## RESULTS

### HNRNPM interacts with SL5 and shuttles into the cytoplasm during SARS-CoV-2 infection

To identify host proteins specifically binding to SL5, we performed pulldowns with biotinylated SL5 RNA (Fig. 1A). A549 cells were chosen as a model cell line for respiratory infection by SARS-CoV-2. Thus, A549 cell lysate was used for fishing of host proteins. As negative controls, we used a shuffled RNA with the same sequence composition as SL5 and the SL5 antisense sequence (Supplementary Table S1). Further, we included a mutated SL5 variant in which all three apical loops in SL5a, SL5b and SL5c, respectively, are mutated to the penta-nucleotide loop 5’-GAGAA-3’. This variant provides a first insight on possible interaction sites of host factors on SL5. Proteins from the pulldown were separated by SDS-PAGE and visualized by Coomassie staining. Comparing the enriched interacting proteins of SL5 and the negative controls, we found two distinct bands representing interacting proteins with a mass of approximately 75 kDa (Fig. 1B). Both, the upper and lower band, were cut out from the SDS gel, the proteins were eluted and subjected to mass spectrometry analysis. In both bands, HNRNPM was identified with the highest Sequest HT score and sequence coverage (Supplementary Table S4 and S5). Thus, both bands were assigned to HNRNPM. This migration pattern of two bands is known for HNRNPM. It represents four different isoforms, with M1 and M2 grouping together in the lower band and M3 and M4 in the upper band (27).

**Fig 1.**
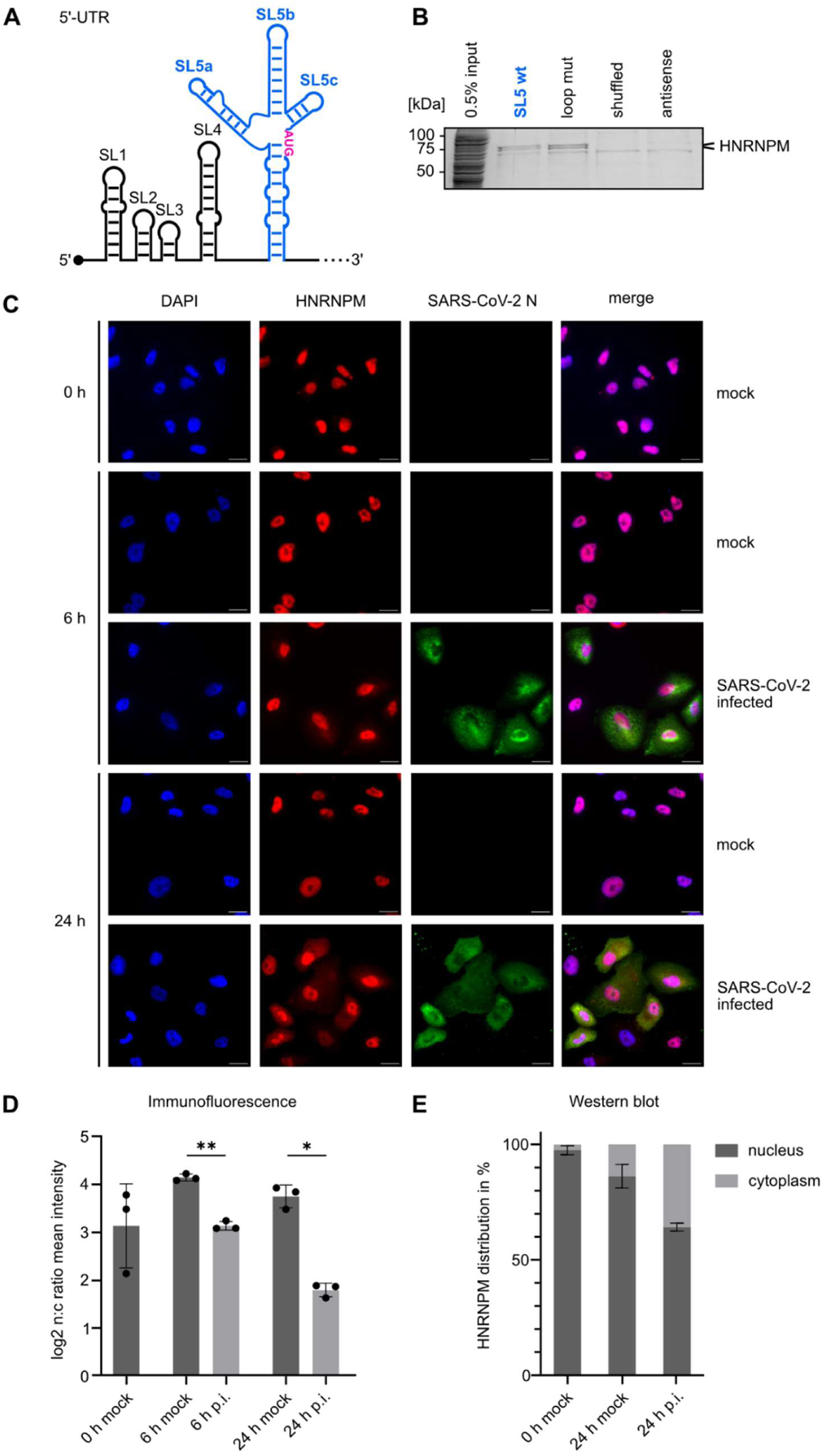
HNRNPM interacts with SL5. (**A**) Schematic representation of the SARS-CoV-2 5’-UTR and first amino acids of nsp1. Stem-loop 5 (SL5) is highlighted in blue. The start codon is highlighted in pink. (**B**) RNA pulldown to identify host proteins in A549 cells interacting with SL5. Proteins from the pulldown were separated by SDS-PAGE and visualized by Coomassie staining. (**C**) Detection of cellular location of HNRNPM with and without infection. A549-hACE2-TMPRSS2 cells were infected with SARS-CoV-2 at an MOI of 0.1. Infected and uninfected control cells were stained with anti-SARS-CoV-2 nucleocapsid antibody (green) and anti-HNRNPM antibody (red). Nuclei were stained with DAPI (blue). Scale bar: 20 µm. n=3. (**D**) Quantification of HNRNPM signal corresponding to (C). 50 cells per condition were quantified. n=3. (**E**) Western blot analysis of cytoplasmic und nuclear HNRNPM fraction of SARS-CoV-2 infected A549-hACE2-TMPRSS2 cells vs. uninfected cells (Supplementary Fig. S1). n=3. Error bars represent standard deviation in all graphs. Statistical significance was assessed by Student’s t-test, two-tailed, paired. *=P<0.05, **= P<0.01. p.i. = post infection

HNRNPM is mainly located in the nucleus (27, 28). Thus, it is unclear how it could directly interact with SL5 on the viral genome, which is located in the cytoplasm. However, it has been shown that HNRNPM shuttles between the nucleus and cytoplasm and enriches in the cytoplasm under certain stress conditions such as viral infections (29, 30). Thus, we determined if HNRNPM also changes its subcellular distribution to the cytoplasm during SARS-CoV-2 infection. For that, we infected A549-hACE2-TMPRSS2 cells with SARS-CoV-2 (BavPat1/2020 isolate) at an MOI of 0.1 and assessed the cellular distribution of HNRNPM by immunofluorescence staining. To verify that SARS-CoV-2 infects A549-hACE2-TMPRSS2 cells, we additionally stained the nucleocapsid protein N in cells. Most of the HNRNPM protein was present in the nucleus regardless of the infection status. However, we observed an increase of cytoplasmic HNRNPM levels in infected cells compared to uninfected cells already within 6 h post infection. At 24 h post infection, cytoplasmic HNRNPM levels were further increased (Fig. 1C). Comparing infected with uninfected cells at 24 h post infection, the log_2_ nuclear/cytoplasmic ratio of the HNRNPM staining was decreased > 2-fold (Fig. 1D).

To further verify shuttling of HNRNPM to the cytoplasm in response to SARS-CoV-2 infection, we performed western blot analysis of subcellular protein fractions from 24 h post infection. Again, successful SARS-CoV-2 infection was verified using an anti-nucleocapsid protein N antibody. As with immunofluorescence, we observed a clear shift in the subcellular distribution of HNRNPM in infected cells compared to uninfected cells with ~ 36% of the HNRNPM protein localized in the cytoplasm (Fig. 1E and Supplementary Fig. S1). Thus, SARS-CoV-2 infection leads to shuttling of HNRNPM into the cytoplasm, where it can interact with the SARS-CoV-2 genome and specifically with SL5.

### HNRNPM promotes protein synthesis from the SARS-CoV-2 5’-UTR

Next, we investigated if the SL5-HNRNPM-interaction affects gene expression by reporter gene assays with the SARS-CoV-2 5’-UTR. To simulate the higher cytoplasmic localization of HNRNPM during SARS-CoV-2 infection, we overexpressed HNRNPM either as full-length protein (HNRNPM FL, ~ 75 kDa) or with disrupted nuclear localization signal (HNRNPM ΔNLS, ~ 69 kDa). eGFP was expressed as control for co-transfection. Western blot analysis verified overexpression of HNRNPM after transfection of plasmids encoding for HNRNPM ΔNLS and HNRNPM FL (Fig. 2A). Analyzing the distribution of HNRNPM between the cytoplasmic and nuclear fraction, we observe ~ 6% of the total HNRNPM protein in the cytoplasmic fraction of the eGFP control. Overexpression of HNRNPM FL leads to a 2-fold increase in the cytoplasmic levels with ~ 12% of total HNRNPM protein. With ~ 22% of HNRNPM in the cytoplasm, HNRNPM ΔNLS expression achieves the highest increase nearly 4-fold over control conditions (Fig. 2B). Thus, expression of HNRNPM ΔNLS most effectively mimicked the HNRNPM translocation after SARS-CoV-2 infection, where we observed ~ 36% cytoplasmic HNRNPM.

**Fig 2.**
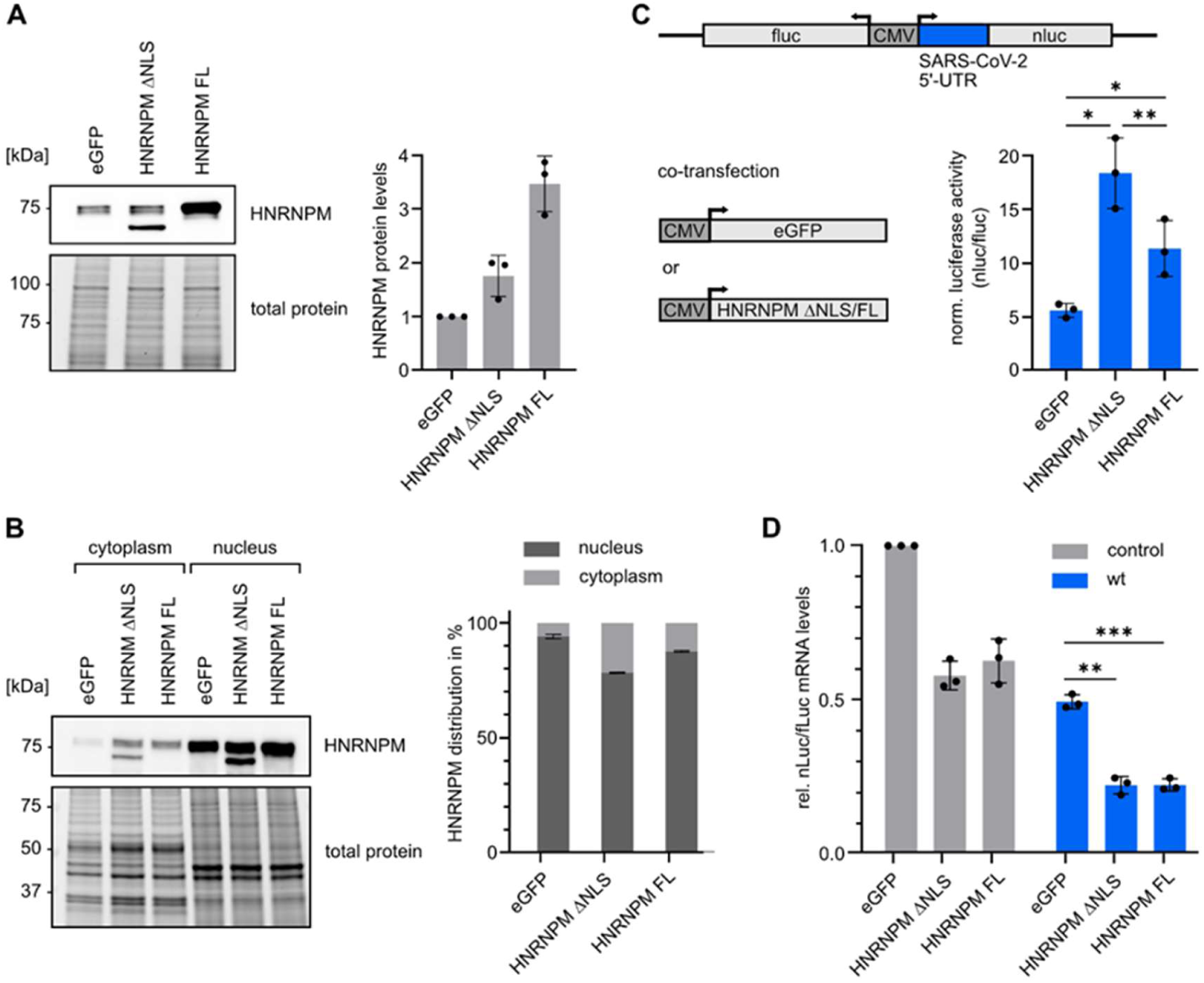
HNRNPM enhances protein synthesis from the SARS-CoV-2 5’-UTR. (A) Western blot of HNRNPM overexpression in A549 cells after transfection of plasmids encoding for eGFP, HNRNPM ΔNLS or HNRNPM FL. Total protein was used as loading control. Protein levels were normalized to an eGFP control. n=3. (B) Western blot to quantify the distribution of HNRNPM between the cytoplasmic and nuclear fractions in A549 cells 24 h after transfection with either a plasmid encoding for eGFP, HNRNPM ΔNLS or HNRNPM FL. Total protein was used as loading control. n=3. (C) Dual luciferase assay to assess the effect of HNRNPM overexpression on protein synthesis. Dual luciferase reporter plasmids with either the SARS-CoV-2 5’-UTR (wt) or empty control were co-transfected with plasmids encoding for eGFP, HNRNPM ΔNLS or HNRNPM FL. Firefly luciferase (fluc) was used as internal control. The ratio of wt nluc/fluc was normalized to the respective ratio of control nluc/fluc. n=3. (D) RT-qPCR to quantify mRNA levels of SARS-CoV-2 5’-UTR (wt) and control after HNRNPM overexpression. Data was normalized to an eGFP control. n=3. Error bars represent standard deviation in all graphs. Statistical significance was assessed by Student’s t-test, two-tailed, paired. *=P<0.05, **= P<0.01, *** =P<0.001. HNRNPM FL = FLAG-tagged full-length HNRNPM, HNRNPM ΔNLS = FLAG-tagged HNRNPM with disrupted nuclear localization signal (Supplementary Table S2).

Since the start codon of the first ORF of SARS-CoV-2 is located on SL5, we investigated the effect of HNRNPM on protein synthesis by the SARS-CoV-2 5’-UTR. We performed dual luciferase assays with firefly luciferase as internal control and NanoLuc luciferase under the control of the SARS-CoV-2 5’-UTR including the sequence encoding the first ten amino acids from nsp1 (nt 1-295) to allow SL5 structure formation (Supplementary Table S3). The reporter was co-expressed with eGFP, HNRNPM ΔNLS or HNRNPM FL. Nluc/fluc ratios were normalized to the nluc/fluc ratios of an empty control vector without SARS-CoV-2 sequences. We observed an ~ 3-fold increased normalized reporter activity with HNRNPM ΔNLS expression compared to the eGFP control and ~ 2-fold increased activity with HNRNPM FL (Fig. 2C). Thus, HNRNPM increases protein production from the SARS-CoV-2 5’-UTR, with a stronger effect for HNRNPM ΔNLS, consistent with the higher amount of cytoplasmic HNRNPM.

To examine if increased protein levels are accompanied by an increase in mRNA levels, we quantified luciferase mRNA levels by RT-qPCR. In contrast to the increased protein level, we observed a reduction in the mRNA level of the SARS-CoV-2 reporter construct by ~ 55% with HNRNPM ΔNLS or FL overexpression compared to the eGFP control. A similar reduction was observed for the mRNA level of the empty control with HNRNPM overexpression compared to eGFP overexpression (Fig. 2D). Thus, this reduction in mRNA levels is an unspecific effect of HNRNPM overexpression and not specific for the SARS-CoV-2 5’-UTR. In sum, HNRNPM positively affects protein production from the SARS-CoV-2 5’-UTR without increasing mRNA levels.

### HNRNPM interaction is mediated by the stem region of SL5

To verify that SL5 binding is necessary for the observed increase in protein synthesis, we performed a mutational analysis. First, we characterized the RNA-protein interaction in more detail by pulldown assays to identify the interacting region of SL5. Pulldown of A549 host proteins interacting with several SL5 variants was performed and enriched proteins were analyzed via western blot with anti-HNRNPM antibody. As the loop mutant with SL5a-c (Fig. 1A and 3A) substituted by 5’-GAGAA-3’ penta-nucleotide loops was still recognized by HNRNPM, we next split SL5 in two halves: the 4-way junction and the stem part of SL5 (Fig. 3A). While the 4-way junction construct was not able to interact with HNRNPM, the stem part retained robust HNRNPM binding. HNRNPM is known to prefer single-stranded RNAs (31). Thus, we designed a stem mutant which preserves the coding region upstream of the start codon (265-294) but removes all single-stranded parts of the stem (ins _157_AAC, _161_AC_162_->G, _168_UAA_170_, del _174_G), forming an uninterrupted duplex. As expected, this double stranded SL5 variant was not able to enrich HNRNPM in pulldowns.

**Fig 3.**
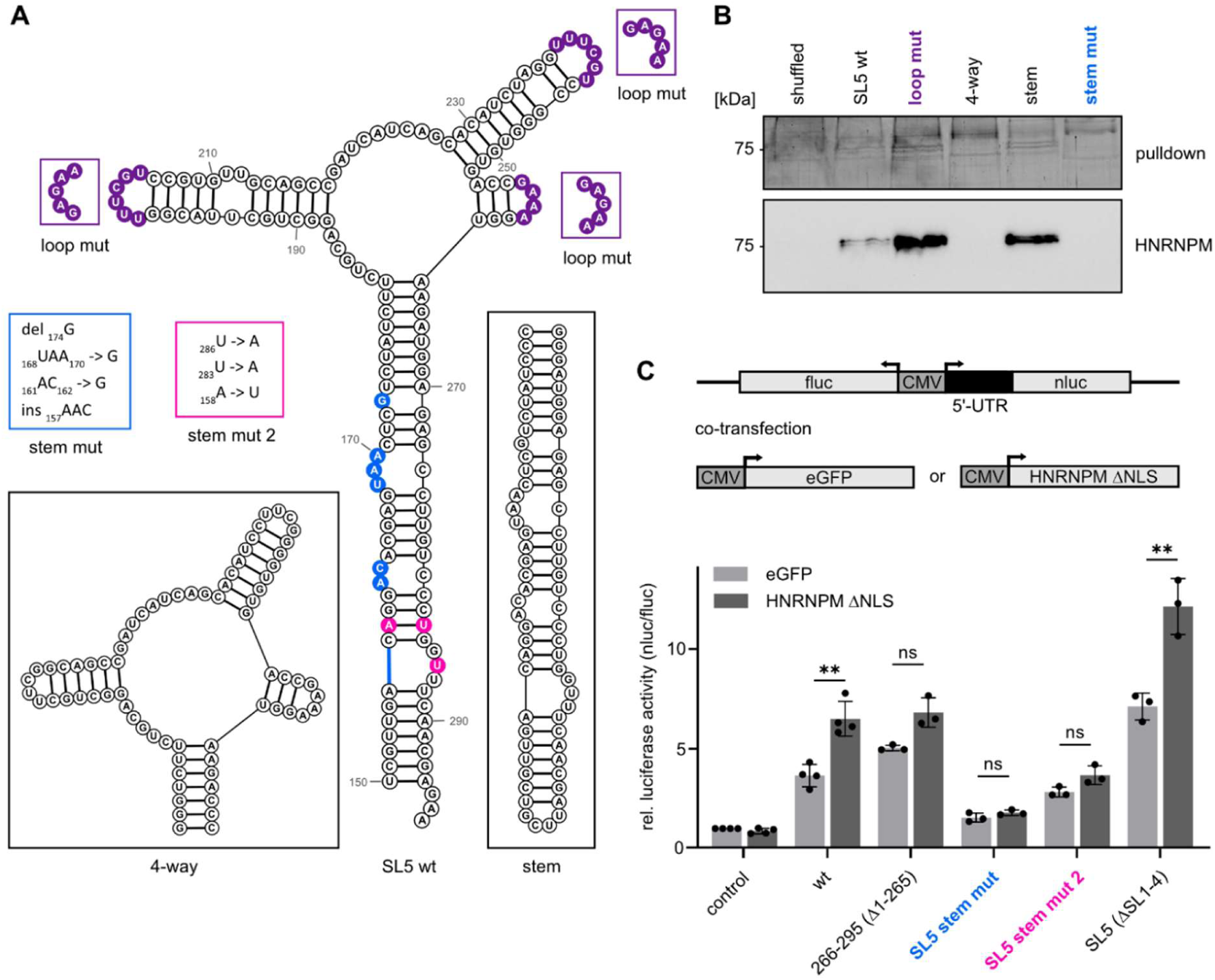
HNRNPM interaction is mediated by the stem region of SL5. (A) Scheme of the secondary structure and sequence of SL5 wt, 4-way, stem, stem mut (changes highlighted in blue), stem mut 2 (changes highlighted in pink) and loop mut (changes highlighted in purple) constructs. (B) SDS-PAGE of A549 host protein pulldown with the indicated SL5 variants. Western blot was performed with anti-HNRNPM antibody. n=3-6. (C) Dual luciferase assay to assess the effect of SL5 mutations or 5’-UTR truncations on reporter activity. Dual luciferase reporter plasmids with NanoLuc luciferase (nluc) as reporter were co-transfected with plasmids encoding for eGFP, HNRNPM ΔNLS or HNRNPM FL. Firefly luciferase (fluc) was used as internal control. Values are normalized to nluc/fluc ratio of empty control reporter plasmid with eGFP overexpression. n=3-4. Error bars represent standard deviation. Statistical significance was assessed by Student’s t-test, two-tailed, paired. **=P<0.01, ns = not significant. HNRNPM ΔNLS = FLAG-tagged HNRNPM with disrupted nuclear localization signal (Supplementary Table S2).

The pulldown data indicate that the stem region of SL5 is responsible for the interaction with HNRNPM. Thus, we tested the SL5 stem mutant in the dual luciferase reporter system to verify that the disrupted binding has an effect on translation efficiency. A respective reporter with the mutated SL5 stem upstream of the NanoLuc luciferase was created (stem mut) and co-expressed with eGFP or HNRNPM ΔNLS (Fig. 3C). Since HNRNPM is described to preferentially bind GU-rich sequences (32), we included a second stem mutant (stem mut 2) with two silent mutations in the coding region (_283_U -> A, _286_U -> A and _158_U -> A). These mutations change the preferential binding motif but preserve the secondary structure of the SL5 stem. Both stem mutants showed reduced luciferase activity and no significant induction upon co-expression of HNRNPM ΔNLS, suggesting that the interaction with the SL5 stem is necessary for modulation of protein synthesis.

Additionally, we constructed a reporter variant, that retains only the sequence upstream of the start codon (266-295) including the GU-rich motif. We observe a slight increase in luciferase activity upon HNRNPM ΔNLS, however not reaching statistical significance. This suggests that the integrity of the SL5 structure is necessary to functionally enhance protein synthesis by HNRNPM. To further verify this finding, we deleted SLs 1-4, keeping only the SL5 stem intact (ΔSL1-4). In this SL5 only construct, luciferase activity was increased compared to the full-length 5’-UTR. This enhancement is likely due to the deletion of SL4, which contains an upstream ORF reducing the usage of the main AUG in SL5 (33). Most importantly, the SL5 only construct showed the same enhancement of luciferase activity upon HNRNPM ΔNLS co-expression as the full-length 5’-UTR.

HNRNPM was previously identified as an IRES *trans*-acting factor (ITAF) (30). In addition, it has been suggested that SL5 can function as part of an IRES (15). This suggests that HNRNPM might upregulate protein synthesis of SARS-CoV-2 by acting as an ITAF. To investigate if a possible IRES activity of the SARS-CoV-2 5’-UTR contributes to HNRNPM regulation, we designed a bicistronic luciferase reporter plasmid with either the HCV IRES, the SARS-CoV-2 5’-UTR or SL5 alone upstream of the NanoLuc luciferase (nluc) ORF (Supplementary Fig. S2A). The reporter plasmid was co-transfected with either a plasmid encoding for eGFP or HNRNPM ΔNLS. Relative luciferase activities of the SARS-CoV-2 SL5 and 5’-UTR bicistronic constructs show only very low luciferase signals compared to the HCV IRES (3-4%). HNRNPM ΔNLS co-expression showed a slight increase in luciferase signals compared to the eGFP controls, but did not result in a significant rescue of the overall very low luciferase activities. Thus, HNRNPM seems not to function as an ITAF for the SARS-CoV-2 5’-UTR in A549 cells.

### RNA recognition motif 1 of HNRNPM is the most essential part for the RNA-protein interaction

HNRNPM comprises three RNA recognition motifs (RRMs) with RRM1 (aa 63-149) and RRM2 (aa 160-242) located as tandem at the N-terminus and RRM3 (aa 606-692) located at the C-terminus, separated by a long flexible linker (Fig. 4A). In order to investigate the domain conferring the SL5 binding preference, we designed several HNRNPM mutants. First, the linker between RRM2 and RRM3 was replaced by a short GGGGS linker (Δ linker) (Fig. 4A), as it has been shown that the linker alone can bind viral RNA (29). FLAG-tagged proteins were overexpressed in A549 cells and RNA pulldown assays were performed with SL5 (Fig. 4A and B). Western blot analysis with anti-FLAG antibody illustrates that full length and Δ linker were equally enriched in the pulldown assay (Fig. 4A and Supplementary Fig. S3A). Thus, the linker seems not to be involved in SL5 binding.

**Fig 4.**
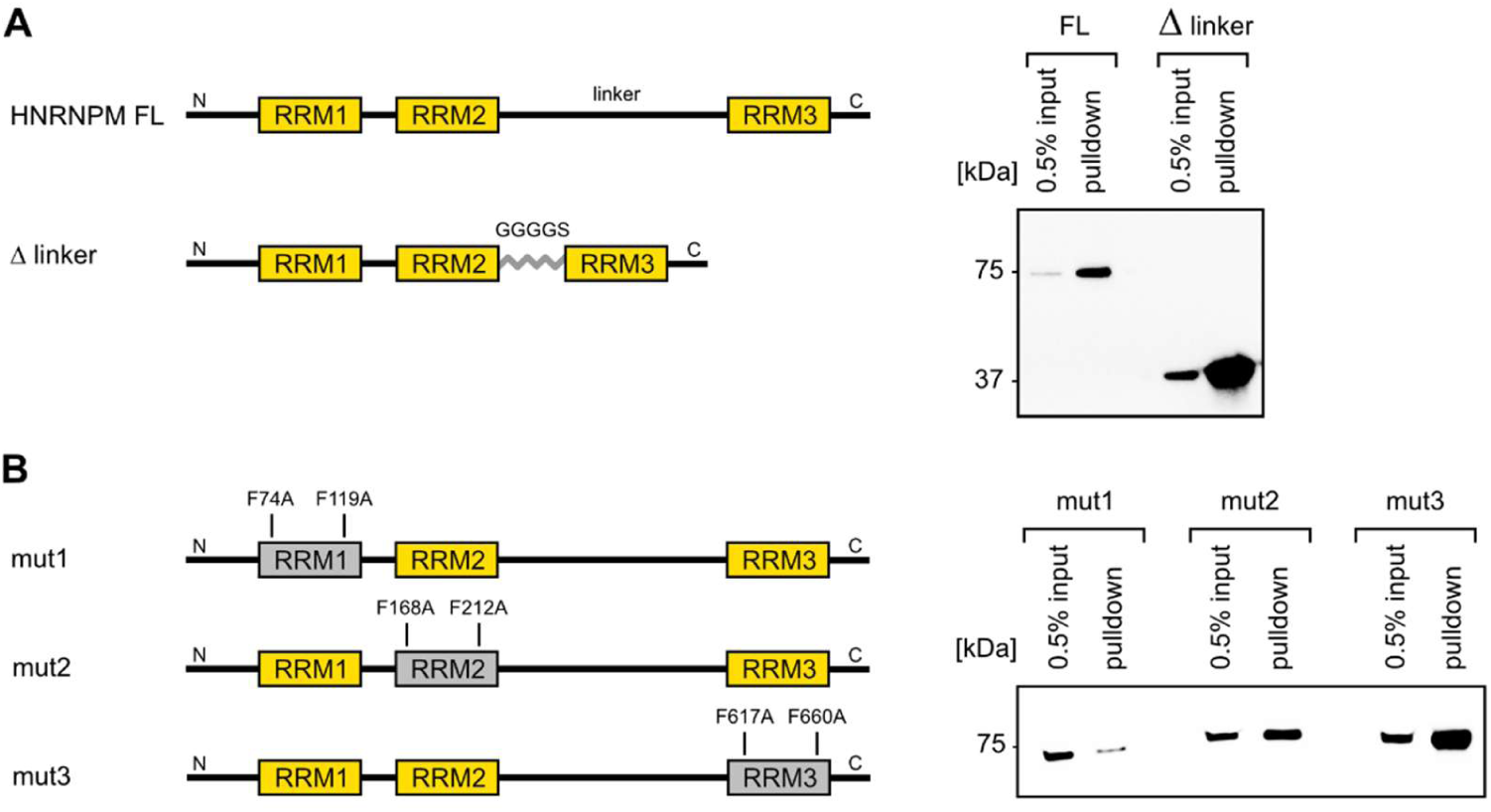
RRM1 is the most important part of HNRNPM for the interaction with SL5. (A) Schematic representation of HNRNPM full-length (FL) and linker mutant (Δ linker). RRMs 1-3 are highlighted in yellow. FLAG-tagged HNRNPM FL and linker mutant expressed in A549 cells were pulled with biotinylated SL5 RNA. Eluted proteins were analyzed via western blot with an anti-FLAG antibody. n=3 (Supplementary Fig. S3A). (B) Overview of HNRNPM RRM mutants (mut1, mut2 and mut3). Two phenylalanine residues in each RRM were mutated as indicated. FLAG-tagged HNRNPM RRM mutants expressed in A549 cells were pulled with biotinylated SL5 RNA. Eluted proteins were analyzed via western blot with an anti-FLAG antibody. n=5 (Supplementary Fig. S3B).

Next, we inspected the individual contribution of the three RRM domains to SL5 recognition. Dependent on the target RNA, each of the three RRMs has been reported to be the most important domain for RNA binding (34–36). Two aromatic residues in the key binding regions (RNP1 and RNP2) mediate RNA recognition in RRM domains (37, 38). Thus, we mutated the respective phenylalanine residues in each RRM (mut1: F74A and F119A, mut 2: F168A and F212A, mut3: F617A and F660A) to obtain three different RRM mutants (Fig. 4B). FLAG-tagged mutant proteins were overexpressed in A549 cells and RNA pulldown assay was performed with SL5 (Fig. 4A and B). Binding of SL5 was most strongly attenuated for mut1 compared to mut2 and mut3, which suggests that RRM1 is crucial for the interaction of HNRNPM with SL5 (Fig. 4B and Supplementary Fig. S3B).

### The interaction between HNRNPM and coronaviral 5’-SL5s is conserved

The higher-order structure of SL5 is highly conserved among different coronaviruses (10, 39). Thus, we examined if HNRNPM can likewise enhance protein synthesis from 5’-UTRs of prominent alpha-(HCoV-229E and HCoV-NL63) and betacoronaviruses (MERS-CoV, MHV-A59 and HCoV-OC43). We constructed dual luciferase reporters with the respective coronaviral 5’-UTRs upstream of the NanoLuc luciferase gene and co-expressed them with either eGFP or HNRNPM ΔNLS. Data were normalized to the luciferase activity of empty control vector. Relative luciferase activity of all coronaviral constructs was increased significantly with HNRNPM overexpression compared to eGFP overexpression (Fig. 5A). In order to confirm the binding of HNRNPM to different SL5s, we performed pulldowns of A549 host proteins with the biotinylated SL5 variants from MERS-CoV and HCoV-229E (Supplementary Fig. S4). SARS-CoV-2 SL5 was included as positive control and antisense RNA of SARS-CoV-2 SL5 as a negative control. The pulldown assay showed the binding of HNRNPM to each tested SL5 variant (Fig. 5B). Thus, HNRNPM might generally act as a proviral factor for coronaviruses by enhancing protein synthesis from the conserved SL5 RNA element.

**Fig 5.**
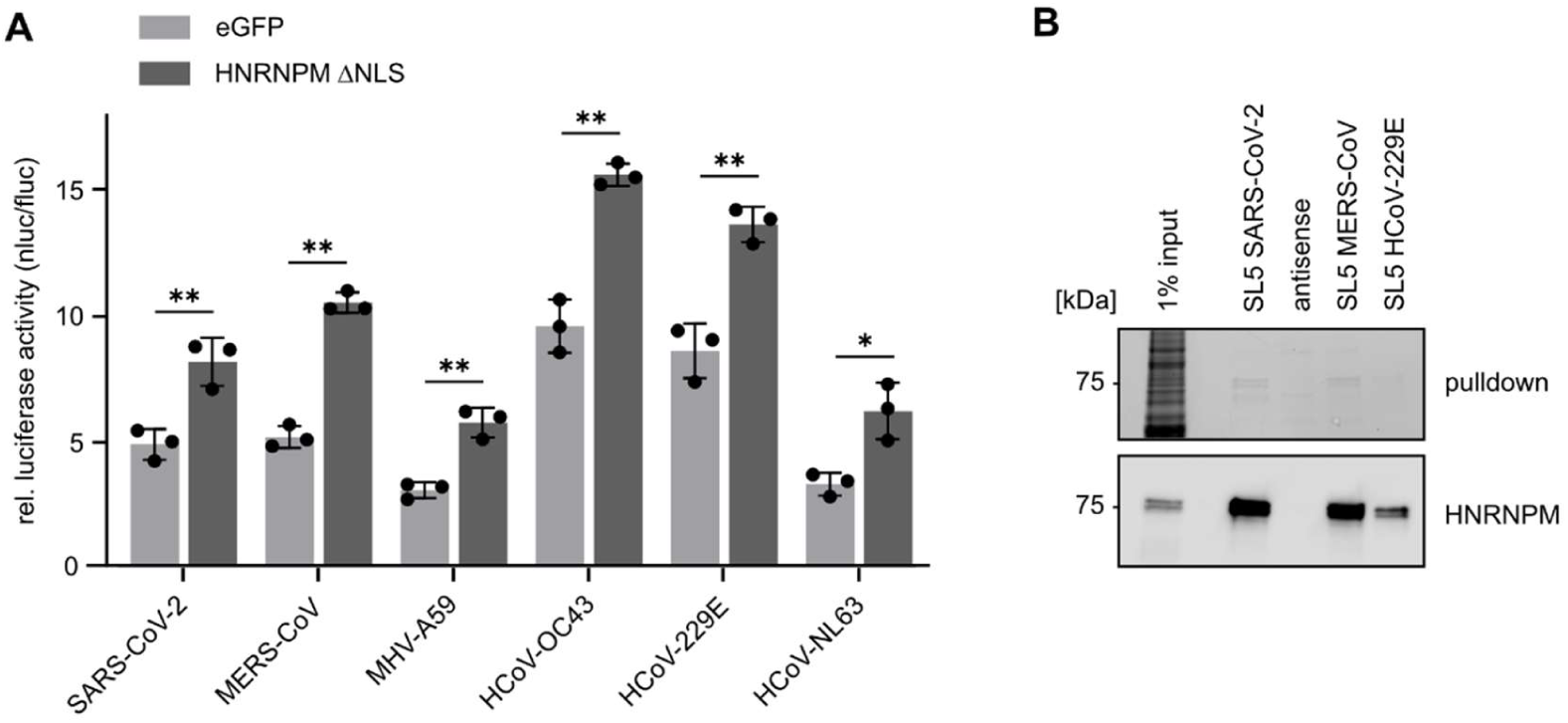
The interaction between HNRNPM and coronaviral 5’-SL5s is conserved. (A) Dual luciferase assay to assess the effect of HNRNPM overexpression on protein synthesis. Dual luciferase reporter plasmids with different coronaviral 5’-UTRs upstream of the NanoLuc luciferase (nluc) ORF were co-transfected with plasmids encoding for eGFP, HNRNPM ΔNLS or HNRNPM FL. Firefly luciferase (fluc) was used as internal control. Values are normalized to nluc/fluc ratio of control reporter plasmid. n=3. Error bars represent standard deviation. Statistical significance was assessed by Student’s t-test, two-tailed, paired. *=P<0.05, **=P<0.01. HNRNPM ΔNLS = FLAG-tagged HNRNPM with disrupted nuclear localization signal (Supplementary Table S2). (B) SDS-PAGE of pulldown with different coronaviral SL5 variants and A549 cell lysate. Western blot was performed with anti-HNRNPM antibody. n=3.

## DISCUSSION

In this study, we identified HNRNPM as an interacting protein of SARS-CoV-2 SL5 and found that HNRNPM shuttles into the cytoplasm during SARS-CoV-2 infection. These observations suggest that HNRNPM can affect the viral life cycle by binding to the SARS-CoV-2 genome. Indeed, we established that cytoplasmic HNRNPM promotes protein synthesis from the viral 5’-UTR by binding to the GU-rich lower stem region of SL5. Comparing two stem mutants (stem mut and stem mut 2), _286_U seems to be particularly involved in the interaction with HNRNPM and is accessible only within a single-stranded sequence. In addition, the structural context is important for the regulation by HNRNPM, as demonstrated with the 266-295 construct, which did not show increased translation efficiency after HNRNPM overexpression to the same extent as folded SL5. Regarding HNRNPM, we found that RRM1 is particularly essential for interacting with SL5. Further, we demonstrated that HNRNPM binds to several SL5s of alpha- and betacoronaviruses and promotes translation to a similar extent, supporting that this mechanism is conserved across different alpha- and betacoronaviruses. Our findings are in line with previous interactome analyses showing that HNRNPM is associated with SARS-CoV-2 RNA (18, 19, 40) and that the knockdown of HNRNPM showed decreased SARS-CoV-2 RNA levels, supporting a proviral function (19). However, the functional role of HNRNPM during infection has not yet been investigated so far. Thus, our observations corroborate earlier evidence that HNRNPM acts as a proviral factor and provide a first understanding of the functional role of HNRNPM.

Further, we provide the first evidence that HNRNPM translocates into the cytoplasm during SARS-CoV-2 infection. Similarly, it has been reported that HNRNPM is redistributed from the nucleus to the cytoplasm during infection with other RNA viruses including Sendai virus (SeV), Encephalomyocarditis virus (EMCV) and Semliki Forest virus (SFV) (29, 41). It has been shown that HNRNPM colocalizes with SFV replication complexes. However, knockdown of HNRNPM increased replication of SFV, indicating an antiviral function in SFV infection (41). Interestingly, HNRNPM has been shown to be a proviral factor for other viruses. During poliovirus and coxsackievirus B3 infection, HNRNPM is cleaved and promotes infection (42). Further, HNRNPM is a proviral factor during SeV and EMCV infection as it reduces antiviral innate immune signaling by binding viral RNA and interacting with RIG-I and MDA5 (29). Notably, SARS-CoV-2 accessory protein ORF6 has been reported to impair the nuclear import of various host proteins by disrupting the Nup98-Rae1-dependent transport (43, 44). Similarly, other hnRNPs have been reported to localize to the cytoplasm upon viral infection e.g., in vesicular stomatitis virus (VSV) infected cells (45). There, knockdown of Rae1 affects cytoplasmic translocation. Whether Rae1 is also required for the translocation of HNRNPM needs to be determined.

HNRNPM is mainly known for its role in pre-mRNA splicing. It associates with intronic sequences in pre-mRNA as part of the pre-spliceosomal A complex (46) and regulates alternative splicing (47). Most often it acts as a splicing repressor, inducing exon skipping or repressing cryptic splice sites (32, 48, 49). Beyond its role in pre-mRNA splicing, HNRNPM has been shown to enhance translation by acting as an ITAF at the *FGF9* IRES under hypoxia (30). Further, it has been reported that HNRNPM interacts with other ITAFs such as Polypyrimidine-tract binding protein (PTB) and PTB-associated splicing factor (PSF) (28). PSF and PTB are characterized as multifunctional RNA-binding proteins involved in splicing and IRES-mediated translation of RNA of several viruses, including coxsackievirus B3 (CVB3) (50), picornavirus (51) and hepatitis C virus (HCV) (52). Already early on, it was suggested that the highly structured 5’-UTR of SARS-CoV-2 may hinder the recruitment of ribosomes to initiate translation and uses a cap-independent mechanism to initiate translation (53). The recently suggested IRES function of SL5, via its conserved apical loops of SL5a and SL5b, has been shown to be most effective under hypoxic conditions (15). Interestingly, HNRNPM also translocates from the nucleus into the cytoplasm under hypoxic conditions (30), suggesting that HNRNPM might support IRES activity. However, we found that the apical loops of SL5a and SL5b are dispensable for binding of HNRNPM to SL5. Further, we found no evidence of IRES-mediated translation of the 5’-UTR of SARS-CoV-2 in A549 cells. It is possible that SARS-CoV-2 requires additional unidentified host co-factors for IRES-dependent translation that are not present at sufficient levels in A549 cells.

Notably, our results are consistent with another study showing exclusively cap-dependent translation initiation and requirement of the eIF4F complex for the SARS-CoV-2 5’-UTR (33). An HNRNPM interactome analysis found proteins involved in splicing and ribosomal function to be strongest enriched (54). Interestingly, the interactome contained several translation initiation factors, including the eIF4F complex components EIF4A1 and EIF4G1. Whether HNRNPM binds ribosome-associated RBPs during viral infection needs to be investigated, as such complexes could promote the recruitment of ribosomes to SARS-CoV-2 RNA.

Without the need for recruitment of other factors, binding of HNRNPM might facilitate start codon recognition simply by destabilizing the RNA structure of SL5. Alternatively, it might enhance start codon recognition by acting as a roadblock that slows down the scanning 43S complex. Such a roadblock mechanism has been shown for RNA structures downstream of start codons and proteins in various eukaryotic systems (55–58). Interestingly, the distance between the AUG and the GU-rich sequence in the SL5 stem is spaced 14 nt apart, which is the distance initially suggested to be optimal to position the 43S complex over the AUG (59). Further work will be needed to determine the precise mechanism of how HNRNPM facilitates translation of the nsps.

A comparative RNA-interactome analysis identified a large overlap between host proteins associated with SARS-CoV-2 and HCoV-OC43 RNA, indicating conserved interactions across betacoronaviruses (19). Notably, HNRNPM was among the host proteins enriched with the RNA of both SARS-CoV-2 and HCoV-OC43. Using computational models trained on CLIP-seq data, another study predicted conserved host RBP binding sites in the 5’-UTR of several coronaviruses and found binding sites for HNRNPM in HCoV-229E and HCoV-NL63 (60). Together, these data support our finding that the HNRNPM-SL5-axis is conserved across human-infecting coronaviruses.

In sum, our data suggest that SL5 is an attractive element for antiviral intervention. Future studies could determine if small molecules can interfere with the binding of HNRNPM to SL5 and attenuate the translation efficiency of the nsps in SARS-CoV-2. The conservation of the interaction between HNRNPM and several coronaviral SL5 further suggests that such interventions could remain effective across several alpha- and betacoronaviruses.

## ACKNOWLEDGEMENTS

This work was supported by the Deutsche Forschungsgemeinschaft (DFG) (project number 495006306 to J.E.W.) and by the LOEWE (State Offensive for the Development of Scientific and Economic Excellence of the German federal state of Hessen) Research Cluster CoroPan (Project 6 to N.B.). We gratefully acknowledge the mass spectrometry core facility at the Department of Chemistry, Marburg University for mass spectrometry analysis.

## DATA AVAILABILITY

The data underlying this study are available in the article and its supplementary material.

